# Eosinophils are dispensable for development of MOG_35-55_-induced experimental autoimmune encephalomyelitis in mice

**DOI:** 10.1101/2021.04.09.439129

**Authors:** Klara Ruppova, Jong-Hyung Lim, Georgia Fodelianaki, Avery August, Ales Neuwirth

## Abstract

Experimental autoimmune encephalomyelitis (EAE) represents the mouse model of multiple sclerosis, a devastating neurological disorder. EAE development and progression involves the infiltration of different immune cells into the brain and spinal cord. However, less is known about a potential role of eosinophil granulocytes for EAE disease pathogenesis. In the present study, we found enhanced eosinophil abundance accompanied by increased concentration of the eosinophil chemoattractant eotaxin-1 in the spinal cord in the course of EAE induced in C57BL/6 mice by immunization with MOG_35-55_ peptide. However, the absence of eosinophils did not affect neuroinflammation, demyelination and clinical development or severity of EAE, as assessed in ΔdblGATA1 eosinophil-deficient mice. Taken together, despite their enhanced abundance in the inflamed spinal cord during disease progression, eosinophils were dispensable for EAE development.

## Introduction

Multiple sclerosis (MS) is an inflammatory autoimmune disorder of the central nervous system (CNS). CNS inflammation by recruited self-reactive T cells and other immune cells, accompanied by activation of microglia, leads to demyelination of neurons and neuroaxonal damage [1]. Various types of myeloid cells infiltrate the CNS and participate in disease pathology [1,2]. Eosinophils are multifunctional granulocytes operating as effector cells in the immune response in different contexts [3]. However, the role of eosinophils for EAE pathogenesis is unclear.

In MS patients, a microarray study revealed the enhanced expression of the gene encoding eosinophil cationic protein (ECP) in active lesions of the brain indicating the occurrence of eosinophils [4]. While eosinophils were only occasionally manifested in the CNS of patients suffering from MS, the eosinophil infiltrates to CNS represent a characteristic hallmark of neuromyelitis optica (NMO) [5]. In the NMO mouse model, eosinophil-derived factors induce damage of astrocytes [6]. Moreover, eosinophils have been observed to infiltrate the spinal cord during EAE [7]. Interestingly, administration of helminth products to EAE mice induced eosinophilia and eosinophil brain infiltration, which was linked with delayed disease onset [8]. However, deficiency of the eosinophil pro-survival factor IL-5 did not affect EAE development [9]. These observations prompted us to investigate a potential role of eosinophils during murine EAE.

## Materials and Methods

### Mice and EAE induction

Wild-type C57BL/6 mice were purchased from Charles River. ΔdblGATA1 mice on C57BL/6 background were previously described [10]. Animal experiments were approved by the Landesdirektion Sachsen, Germany.

MOG_35-55_ peptide (myelin oligodendrocyte glycoprotein; peptides&elephants) was mixed with Freund’s adjuvant containing 500 μg of inactivated *Mycobacterium tuberculosis* H37RA (Difco). After emulsification, the peptide (200 µg) was subcutaneously injected to female mice (8-12 weeks old). Subsequently, mice received via i.p. 400 ng of pertussis toxin (Merck) which was repeated two days later. Clinical score was determined as follows: 0=no clinical sign; 1=limp tail; 2=mild hind limb weakness; 3=complete unilateral hind limb paralysis or partial bilateral hind limb paralysis; 4=complete bilateral hind limb paralysis; 5=four limb paralysis; 6=death. EAE mice were analyzed at pre-onset (day 7 post immunization), onset (day 9-13 post immunization) and peak (day 17-19 post immunization) of the disease.

### Flow cytometry analysis

Spinal cord was collected from euthanized mice after systemic perfusion and Myelin Removal Beads (Miltenyi Biotec) were applied to obtained leukocyte-enriched fraction. Following antibodies were used: anti-CD45-PE, anti-CD11b-AF488 and anti-Siglec-F-AF647 (BD Biosciences); anti-CCR3-PE-Vio770, anti-CD4-APC, anti-CD8-PerCP (Miltenyi Biotec); anti-Ly6G-PerCP/Cy5.5 (BioLegend); anti-F4/80-PE/Cy7 (eBioscience). FACSCanto II and FlowJo software were used for analysis.

### Real-time quantitative PCR (RT-qPCR)

RNA was transcribed to cDNA using iScript (Bio-Rad). RT-qPCR was performed using SsoFast EvaGreen Supermix (Bio-Rad). Gene expression was analyzed by ΔΔCT method with RPS29 as reference gene. Sequences of primers are available upon request.

### Eotaxin-1 ELISA and cytokine measurement

Tissues were collected in PBS supplied with protease inhibitors (Roche). Samples were homogenized, centrifuged and supernatants were analyzed using Mouse CCL11/Eotaxin Quantikine ELISA (R&D). To measure cytokines, samples were homogenized in RIPA buffer supplied with protease inhibitors. V-PLEX Proinflammatory Panel 1 kit was used for cytokine measurement (Meso Scale Discovery). Concentration of eotaxin-1 and cytokines was normalized to the total protein amount assessed by BCA (Thermo Fisher Scientific).

### Luxol fast blue staining

Spinal cord was fixed with 4% paraformaldehyde, embedded in OCT (Sakura) and cut to 12 μm sections that were stained with Luxol Fast Blue (LFB) stain (Sigma). Mosaic images were acquired using a Zeiss Axio Observer.Z1-Inverted Microscope with Apotome. Four to six sections from each mouse were analyzed using ImageJ. Myelin concentration was determined as integrated density after automatic thresholding and divided by the section area. To normalize, values of WT mice were set as 100 %.

### Statistical analysis

Mann Whitney test was calculated with GraphPad6 software. P values ≤0.05 were considered statistically significant. Data are presented as mean+SEM.

## Results

### Eosinophil abundance in the spinal cord increases during EAE

To examine the role of eosinophils in EAE, we first monitored eosinophil numbers in the inflamed spinal cord by flow cytometry; eosinophils were characterized as CD45+SSC^high^FSClow CD11b+Siglec-F+ cells (Figure 1A). In comparison to healthy mice, there was an increase in eosinophil numbers at onset and peak of EAE (Figure 1B). Eosinophil accumulation was verified by mRNA analyses (RT-qPCR) of eosinophil-specific granular proteins (ECP, PRG2) (Fig. 1C-D). In addition, eotaxin-1 protein was elevated in the spinal cord during the course of EAE (Fig. 1E) indicating a possible mechanism for active eosinophil recruitment.

**Figure 1.**
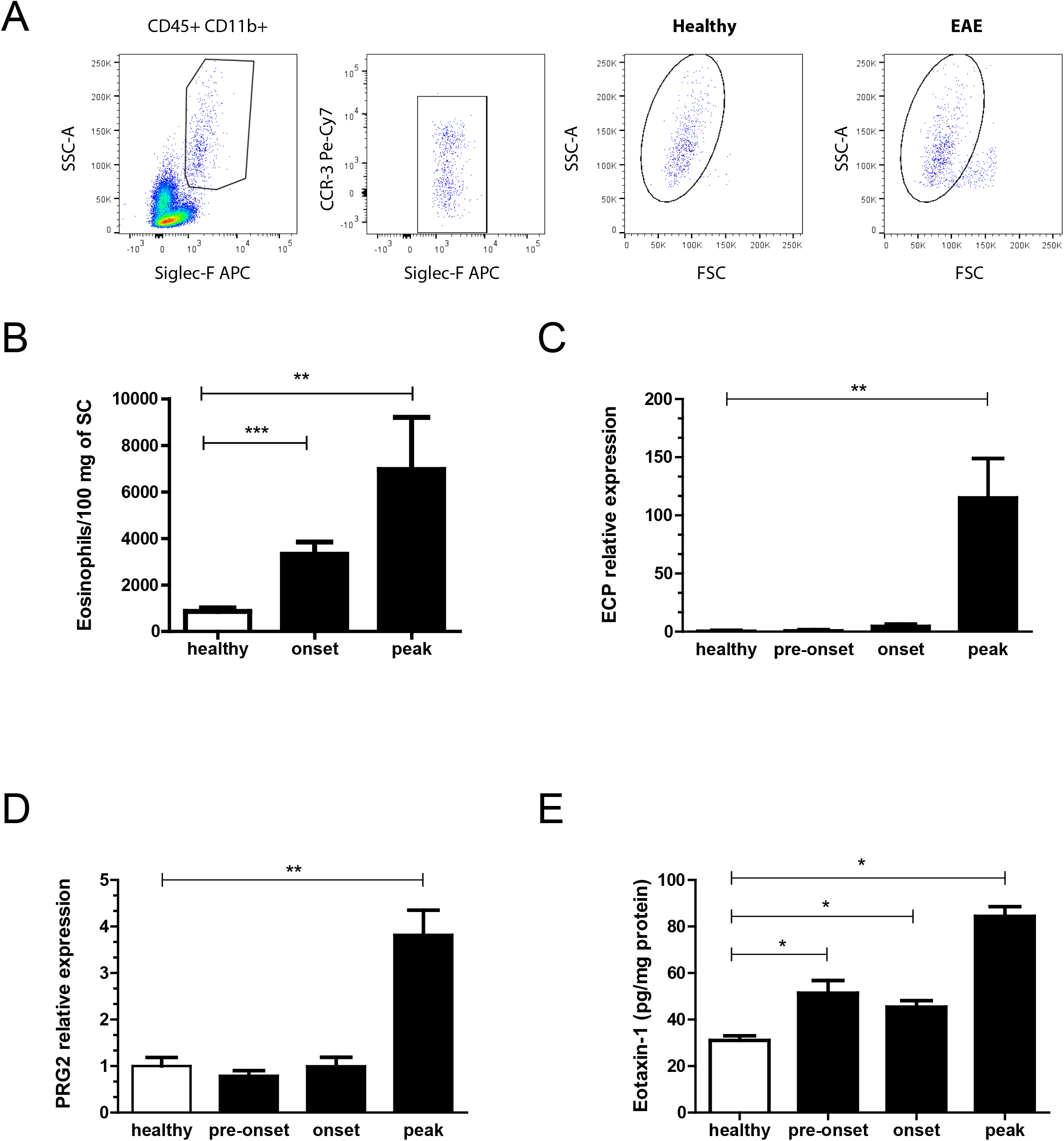
Eosinophil abundance in the spinal cord increases in the course of EAE. **(A)** Flow cytometry analysis for eosinophils in the spinal cord of healthy and EAE mice; eosinophils were identified as CD45+ SSChigh FSClow CD11b+ Siglec-F+ cells. Representative plots are shown to depict the gating strategy. Numbers of **(B)** eosinophils present in the spinal cord of healthy mice or EAE mice at onset and peak of the disease (n = 9-10/group). Relative mRNA expression of **(C)** ECP and **(D)** PRG2 in the spinal cord of healthy mice or EAE mice at pre-onset, onset and peak of the disease (n = 6-7/group). Relative mRNA expression in the spinal cord of healthy mice was set as 1. **(E)** Protein levels of eotaxin-1 analyzed by ELISA in spinal cord homogenates from healthy and EAE mice at pre-onset, onset and peak of the disease. Eotaxin-1 concentration was normalized to the total protein amount (n = 4-5/group). * P≤0.05, ** P≤0.01. Data are presented as mean ± SEM. ECP (eosinophil cationic protein), PRG2 (major basic protein), SC (spinal cord). Data are representative of two independent experiments (B).

### Eosinophils have no impact on EAE

Since we found that eosinophils accumulate in the spinal cord during EAE, we next assessed their contribution to disease development using eosinophil-deficient ΔdblGATA1 mice [11]. No significant difference in disease severity between eosinophil deficient and control ΔdblGATA1 mice was observed (Fig. 2A). In accordance, we did not find any significant changes in the numbers of infiltrating cells except of eosinophils at the peak of EAE using flow cytometry analyses (Fig. 2B, C). Moreover, demyelination, assessed histochemically by LFB staining for myelin, and spinal cord inflammation, assessed by mRNA expression analysis of inflammatory molecules followed by multiplex cytokine measurement, was not different between ΔdblGATA1 and WT mice (Fig. 2D-H). Expectedly, expression of the eosinophil marker PRG2 was strongly reduced in ΔdblGATA1 mice (Fig. 2D). Together, EAE disease severity, neuroinflammation and demyelination were not affected by eosinophil deficiency.

**Figure 2.**
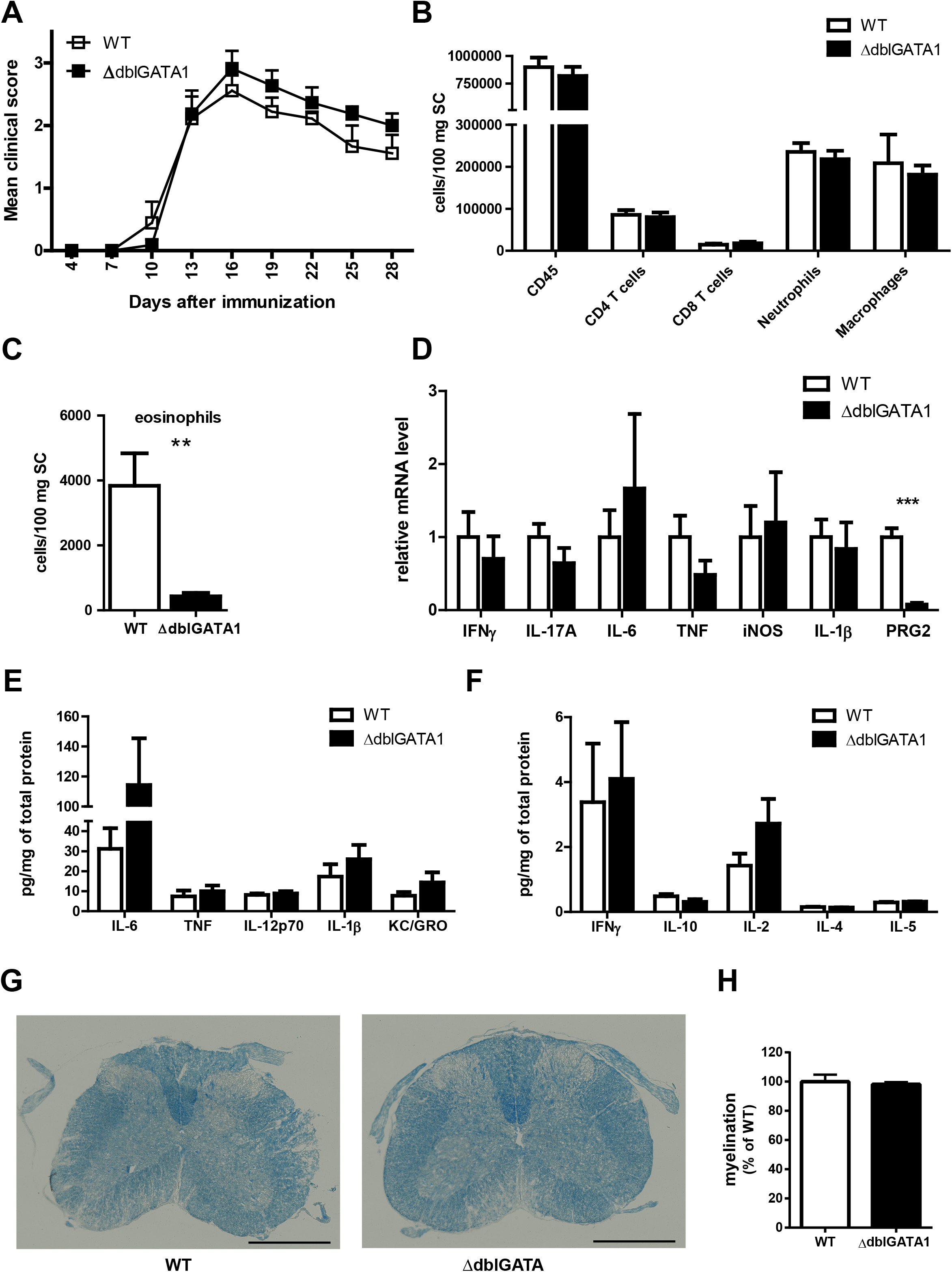
Eosinophils are dispensable for EAE development. **(A)** EAE was induced in ΔdblGATA1 mice and their WT controls and the clinical development was scored (n=9-11/group). Mean clinical score was plotted in a time course. **(B-H)** EAE was induced in ΔdblGATA1 mice and their WT controls and the spinal cord was analyzed at the peak of the disease. **(B)** Cell numbers of total CD45+ leukocytes, CD4 T cells (CD45+ CD8-CD4+), CD8 T cells (CD45+ CD4-CD8+), neutrophils (CD45+ CD11b+ F4/80-Ly6G+), macrophages (CD11b+ Ly6G-F4/80+ CD45high) and **(C)** eosinophils analyzed by flow cytometry (n=4-6/group). **(D)** Expression of IFNγ, IL-17A, IL-6, TNF, iNOS, IL-1β and PRG2 mRNA in the spinal cord was analyzed using RT-qPCR (n=7/group). Gene expression of WT mice was set as 1. **(E-F)** Protein levels of cytokines analyzed by multiplex assay. Cytokine concentration was normalized to the total protein amount (n=4-5/group). **(G)** Spinal cord cryosections were stained with LFB for assessment of myelin. Representative images depicting LFB staining in eosinophil-deficient and WT mice. Scale bar 500 μm. **(H)** Quantification of myelination is shown relative to WT mice (n=3-5/group). ** P≤0.01, *** P≤0.001. Data are presented as mean ± SEM. SC (spinal cord), LFB (Luxol Fast Blue). Data are representative of two independent experiments (A-C).

## Discussion

Although it has been previously reported that eosinophils infiltrate into the CNS during EAE, their impact on disease pathology remained controversial [7-9]. Here, we found increased eosinophil abundance in the spinal cord during EAE, which might be attributed at least in part to enhanced levels of eotaxin-1 in inflamed spinal cord. Although this observation resulted in the hypothesis that eosinophils may influence EAE pathogenesis, our data utilizing eosinophil-deficient mice revealed that eosinophils are dispensable for EAE development and progression. Presumably, eosinophil role may be only minor or other more abundant immune cells in the inflamed spinal cord may compensate for the absence of eosinophils. Contrary to our study, injection of helminth products leads to IL33-dependent eosinophilia together with delayed onset and development of EAE. [8]. However, in the context of helminth infection, eosinophils may acquire a regulatory phenotype, thereby resulting in EAE amelioration. Indeed, adoptive transfer of *in vitro* differentiated eosinophils from the bone marrow did not reduce EAE severity. Moreover, helminth products may induce additional immunomodulatory mechanisms, such as regulatory T cells or regulatory cytokines [12-14]. On the contrary, adoptive transfer of eosinophils from interleukin-33 treated mice delay onset and decreased severity of EAE [8]. Alongside the concept that eosinophils bearing different phenotypes have diverse roles in immunity, Mesnil *et al*. described in asthmatic lungs not only the population of pathogenic pro-inflammatory eosinophils but characterized tissue resident eosinophils with opposed, regulatory function [15]. Our data together with these previous studies indicate that only eosinophils shifted to a specific phenotype (e.g. in the context of a parasitic infection) may be capable of modulating the immune response during EAE development.

Taken together, we demonstrated that eosinophils accumulate in the inflamed spinal cord in the course of EAE. However, eosinophils have no impact on the development and severity of EAE at least in our experimental setting, i.e. in EAE induced in C57BL/6 mice by immunization with MOG_35-55_ peptide.

## Abbreviations

MS: Multiple sclerosis
EAE: experimental autoimmune encephalomyelitis
CNS: Central nervous system
ECP: Eosinophil cationic protein
IL-5: Interleukin-5
IL-33: Interleukin 33
NMO: Neuromyelitis optica
LFB: Luxol Fast Blue
MOG: myelin oligodendrocyte glycoprotein
IFNγ: interferon gamma
IL-17A: interleukin 17A
IL-6: Interleukin-6
TNF: tumour necrosis factor
iNOS: inducible nitric oxide synthase
IL-1β: interleukin 1 beta
PRG2: eosinophil major basic protein
SC: spinal cord.

## Acknowledgement

We thank T. Chavakis (Technische Universität Dresden, Germany) for helpful discussions and S. Grossklaus (Technische Universität Dresden, Germany) for technical assistance.

## Author Contributions

KR designed and performed the experiments, analyzed and interpreted data and wrote the manuscript. JL and GF performed experiments and analyzed data. AA provided ΔdblGATA1 mice and edited the manuscript. AN designed and performed experiments, analyzed and interpreted the data and wrote the manuscript.

## Data availability

The data are available from the corresponding author upon request.

